# Channel nuclear pore protein 54 directs sexual differentiation and neuronal wiring required for female reproductive behaviors in *Drosophila*

**DOI:** 10.1101/806380

**Authors:** Mohanakarthik P. Nallasivan, Irmgard U. Haussmann, Alberto Civetta, Matthias Soller

**Author notes:** equal contributing authors (Mohanakarthik P. Nallasivan, Irmgard U. Haussmann).

## Abstract

The post-mating response induced by male-derived sex-peptide in *Drosophila* females is a well-established model to elucidate how complex innate behaviors are hard-wired into the brain. Here, we found that the channel nuclear pore protein *Nup54* gene is essential for the sex-peptide response as viable mutant alleles do not lay eggs and reduce receptivity upon sex-peptide exposure. Nup54 directs correct wiring of few adult brain neurons that express *pickpocket* and are required for egg laying, but channel Nups also mediate sexual differentiation and male X-chromosome dosage compensation. Consistent with links of Nups to speciation, the Nup54 promoter is a hot spot for rapid evolution and promoter variants alter expression in transgenes. These results implicate altered expression of Nup54 to the onset of speciation processes leading to changes in neuronal wiring and sexual differentiation as a response to sexual conflict arising from male-derived SP to direct the female post-mating response.

## Introduction

Female reproductive behaviors and physiology change profoundly after mating. Control of pregnancy-associated changes in physiology and behaviors are largely hard-wired into the brain to guarantee reproductive success, yet the gene expression programs that direct neuronal differentiation and wiring at the end of the sex determination pathway are largely unknown.

In most insects, male-derived substances transferred during mating direct female physiology and post-mating behaviors [1-3]. *Drosophila* females display a repertoire of sex specific behaviors after mating including reduced receptivity (readiness to mate) and increased egg laying [4, 5]. The main trigger of these post-mating behaviors is male-derived sex-peptide (SP) [6-9]. SP is a 36 amino acid peptide that is transferred during mating to the female [10]. Besides reducing receptivity and increasing egg laying, SP induces a number of other behavioral and physiological changes including increased egg production, feeding, a change in food choice, sleep, constipation and stimulation of the immune system [11-18]. In addition, SP binds to sperm and acts as a sperm sensor, is required for the release of stored sperm and imposes costs of mating [19-21].

Many of the phenotypic effects of SP on female physiology are known, but central aspects of the neuronal circuitry governing regulation of the main post-mating behaviors like reduced receptivity and increased oviposition are unclear. First insights came from the analysis of *egghead* mutant alleles that are insensitive to SP [22]. In these mutants neuronal connections from the ventral nerve cord to the central brain show defects resulting in an egg retainer phenotype and non-responsiveness to SP in reducing receptivity. This suggests that receptor signaling is disconnected from the motor output programs.

Other attempts to map the circuitry mediating the post-mating response used expression of membrane-tethered SP or RNAi knockdown of a receptor for SP, SPR, in distinct neuronal expression patters [9, 23-27]. These screens identified *pickpocket* (*ppk*), *fruitless* (*fru*) and *dsx* expressing neurons mediating the post-mating switch via expression of membrane-tethered SP in multiple pathways [9].

The ability of males to manipulate post-mating responses of females, for example, by SP in *Drosophila*, can promote sexual conflict and trigger an arms-race between the sexes [28-30]. This arms-race drives rapid evolution and can fuel speciation. Few genes have been linked to speciation by mapping inviability or sterility in hybrids of close relatives. Among them are two essential nuclear pore proteins, *Nup96* and *Nup160*, that are part of the outer ring of the pore and interact with each other trigger inviability in species hybrids through adaptive divergence in these two proteins [31-34]. The megadalton nuclear pore complex constitutes a bidirectional gateway connecting the nucleus and cytoplasm to control transport of all macromolecules [35], but how nuclear pore proteins can drive speciation is unknown.

Here, we have identified a role for the essential *Nuclear pore protein 54* (*Nup54*) gene, that is part of the nucleo-cytoplasmic transport channel, in constituting the SP response through an allele displaying egg retention and insensitivity to SP with regard to receptivity. Nup54 is a core protein of the nuclear pore and localizes to the transport channel connecting the nucleus with the cytoplasm. Nup54 is required early in neuronal development to direct proper wiring of few neurons in the adult brain that express *ppk* and are part of the neuronal circuit directing egg laying. In addition, more general roles for the nuclear pore in regulating sexual differentiation are indicated by RNAi knockdown of channel Nups, which results in differentiation defects of internal and external sexual features in males and females. Consistent with a role in sexual conflict driven speciation, the *Nup54* promoter is a hot-spot for rapid diversification and the identified *Nup54*^*QB62*^ promoter deletion allele as well as the promoter from closely related *D. simulans* alter expression of transgenes.

## Results

### Mutant females for *Nup54* alleles display an egg retainer phenotype and are insensitive to sex-peptide for reducing receptivity

To identify novel genes involved in specifying the SP response, we screened females from egg retainer lines that have normal oogenesis, but are unable to lay eggs, for their ability to reduce receptivity [22]. Using this approach, we identified a homozygous viable EMS induced line, *QB62* [36], that did not reduce receptivity upon injection of SP (Figure 1A-D). We mapped this allele by meiotic recombination based on the egg retention phenotype to 2-69.9, corresponding approximatively to chromosome position 50C. Using overlapping deficiencies this allele could then be mapped to chromosome section 49A4 by deficiencies *Df(2R)BSC305* and *Df(2R)Exel16061* (Figure 1A). To further restrict the number of genes in this chromosomal area we generated a smaller deficiency, *Df(2R)9B4*, by FRT mediated recombination between transposons *P{XP}CG8525*^*d06853*^ and *PBac{RB}DUBAI*^*e00699*^ (Figure 1A).

**Figure 1:**
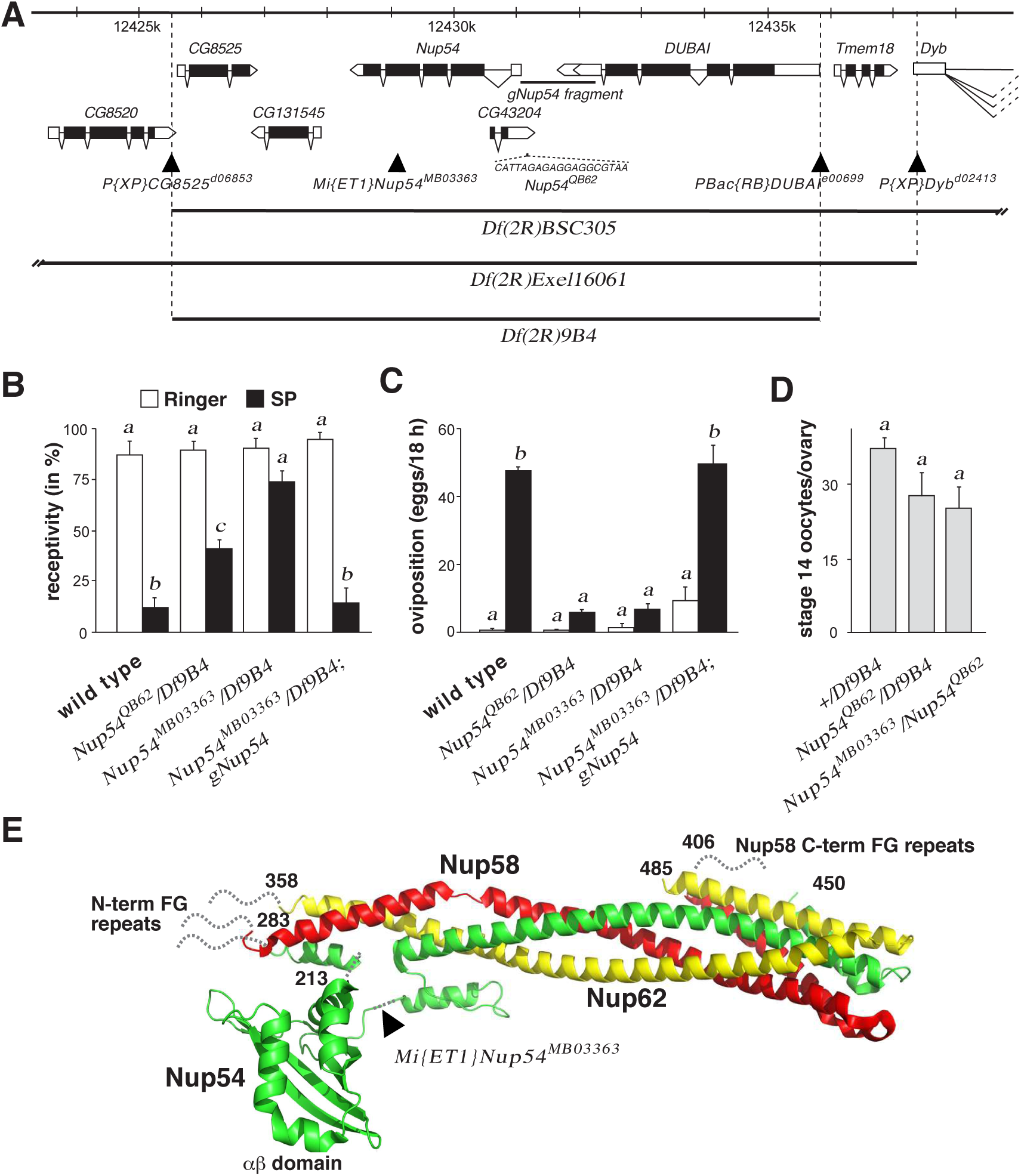
Mapping of sex-peptide insensitive EMS allele *QB62* to *Nup54*. **A)** Schematic of the *Nup54* chromosomal region depicting gene models and chromosomal deficiencies used below the chromosomal nucleotide positions. Coding parts are shown as black and non-coding parts as white boxes. Transposon insert are show as triangles. The sequence of the deletion in *Nup54*^*QB62*^ and the promoter fragment used for genomic rescue construct *gNup54* are shown below the gene model. **B)** Receptivity of wild type and transheterozygous *Nup54*^*QB62*^*/Df(2R)9B4, Nup54*^*MB03363*^*/Df(2R)9B4* and *Nup54*^*MB03363*^*/Df(2R)9B4; gNup54* females after sex-peptide (SP, black) or Ringer’s (R, white) injection measured by counting mating females in a 1 h time period 3 h after SP or R injection, respectively. Means with the standard error for three experiments with 18-23 females each are shown, and statistically significant differences are indicated by different letters (p≤0.001). **C)** Oviposition of wild type and transheterozygous *Nup54*^*QB62*^*/Df(2R)9B4, Nup54*^*MB03363*^*/Df(2R)9B4* and *Nup54*^*MB03363*^*/Df(2R)9B4; gNup54* females after sex-peptide (SP, black) or Ringer’s (R, white) injection shown as means of eggs laid in 18 h with the standard error for 10 females each, respectively, and statistically significant differences are indicated by different letters (p≤0.001). **D)** Number of stage 14 oocytes present in ovaries of sexually mature virgin wild type and transheterozygous *Nup54*^*QB62*^*/Df(2R)9B4* and *Nup54*^*MB03363*^*/Df(2R)9B4* females shown as means with the standard error for six ovaries. **E)** Ribbon diagram of the structure of the channel Nup complex consisting of Nup54, Nup58 and Nup62 with the position of the *Mi{ET1}Nup54*^*MB03363*^ transposon indicated.

We then identified a *Mi{ET1}*transposon insert in the *Nup54* gene, *Nup54*^*MB03363*^, that was allelic to *QB62* with respect to both egg retention and reduction of receptivity after SP injection (Figure 1B-D). Nup54 localizes to the transport channel of the megadalton nuclear pore complex that constitutes a bidirectional gateway connecting the nucleus and cytoplasm to control transport of all macromolecules [35]. The *Nup54*^*MB03363*^ transposon insert leads to a truncation of the Nup54 ORF and removes the C-terminal alpha helical domain that connects Nup54 with Nup58, but leaves the core part containing the FG repeats composing the inner nuclear channel and the interaction of Nup54 with Nup62 via the alpha/beta helical region intact (Figure 1E) [35, 37, 38]. Sequencing of the *Nup54*^*QB62*^ allele identified a small deletion (CATTAGAGAGGAGGCGTAA) in the promoter region 186 nt upstream of the first transcribed nucleotide. These two alleles represent hypomorphic mutations as channel Nups are essential genes [35].

A genomic rescue using a construct with a deletion of the first intron containing the overlapping gene *CG43204* (*gNup54*) further confirmed that the egg retention and SP insensitivity in receptivity maps to the *Nup54* gene (Figure 1B and C).

We then analyzed the expression of Nup54 from *gNup54* via the C-terminal HA tag. Nup54 localizes to the nuclear membrane in salivary glands (Fig S1A) and is broadly expressed in the brain. Intriguingly, when Nup54 is expressed from *gNup54*^*QB62*^ harboring the promoter of the *QB62* allele, expression is much reduced in salivary glands and altered in the brain (Fig S1A-D).

On a gross level, brain morphology looks normal in *Df(2R)9B4 gNup54*^*QB62*^ (Fig S1B, D) arguing for more specific defects in the brain. To evaluate brain morphology in more detail we analysed neuronal projections of *dsx* neurons, as these neurons are directly relevant to female sexual behaviors. In both the ventral nerve cord and the brain we did not detect alterations of *dsx* neuronal projections *Df(2R)9B4 gNup54*^*QB62*^ compared to wild type (Fig S1 G-J).

### Nup54 is required before neuronal maturation for establishing the post-mating response

Next, we expressed *Nup54* from a *UAS* construct in a *Nup54*^*MB03363*^*/Df(2R)9B4* background to identify its temporal and spatial requirement for rescuing the egg retainer and SP insensitivity phenotype in receptivity. Global expression with *tubGAL4* in *Nup54*^*MB03363*^*/Df(2R)9B4* completely rescued these phenotypes after injection of SP. We then expressed *UASNup54* in all differentiated neurons with *elavGAL4*^*C155*^ or in subsets using *ppkGAL4, fruGAL4* and *dsxGAL4* to test whether they can rescue the post-mating response in *Nup54*^*MB03363*^*/Df(2R)9B4* after SP injection (Figure S2A and B). These experiments revealed that *fruGAL4* and *dsxGAL4*, which express earlier than *elav*, can rescue the SP insensitivity and suppress remating, but only dsxGAL4 can marginally rescue egg laying (Figure S2A and B).

Taken together these data indicate that Nup54 is required early in development before neurons are fully differentiated to specify the neuronal circuits required for the post-mating response and that the neuronal circuits for egg laying and receptivity are specified by distinct neuronal populations.

### *Nup54*^*QB62*^ displays defects in neuronal wiring of *pickpocket* expressing neurons

Expression of membrane-bound SP can induce the post-mating response in *ppk* expressing neurons [23-26]. To examine whether Nup54 function is required for neuronal wiring relevant for the post-mating response, we visualized neuronal projections of *ppk* neurons in wild type and *Nup54*^*QB62*^*/Df(2R)9B4* mutant females by expression of CD8::GFP from *UAS* under the control of *ppkGAL4*. In the larval brain of control females *ppk* sensory neuronal projections display a regular pattern of connectives and commissures, while in *Nup54*^*QB62*^*/Df(2R)9B4* mutants many connectives and commissures are disrupted (Figure 2A-F). In the adult brain, projections in the central brain were dramatically reduced in *Nup54*^*QB62*^*/Df(2R)9B4* females compared to control females (Figure 2G and H), while no projection differences were obvious in the ventral nerve cord (Figure J and K).

**Figure 2:**
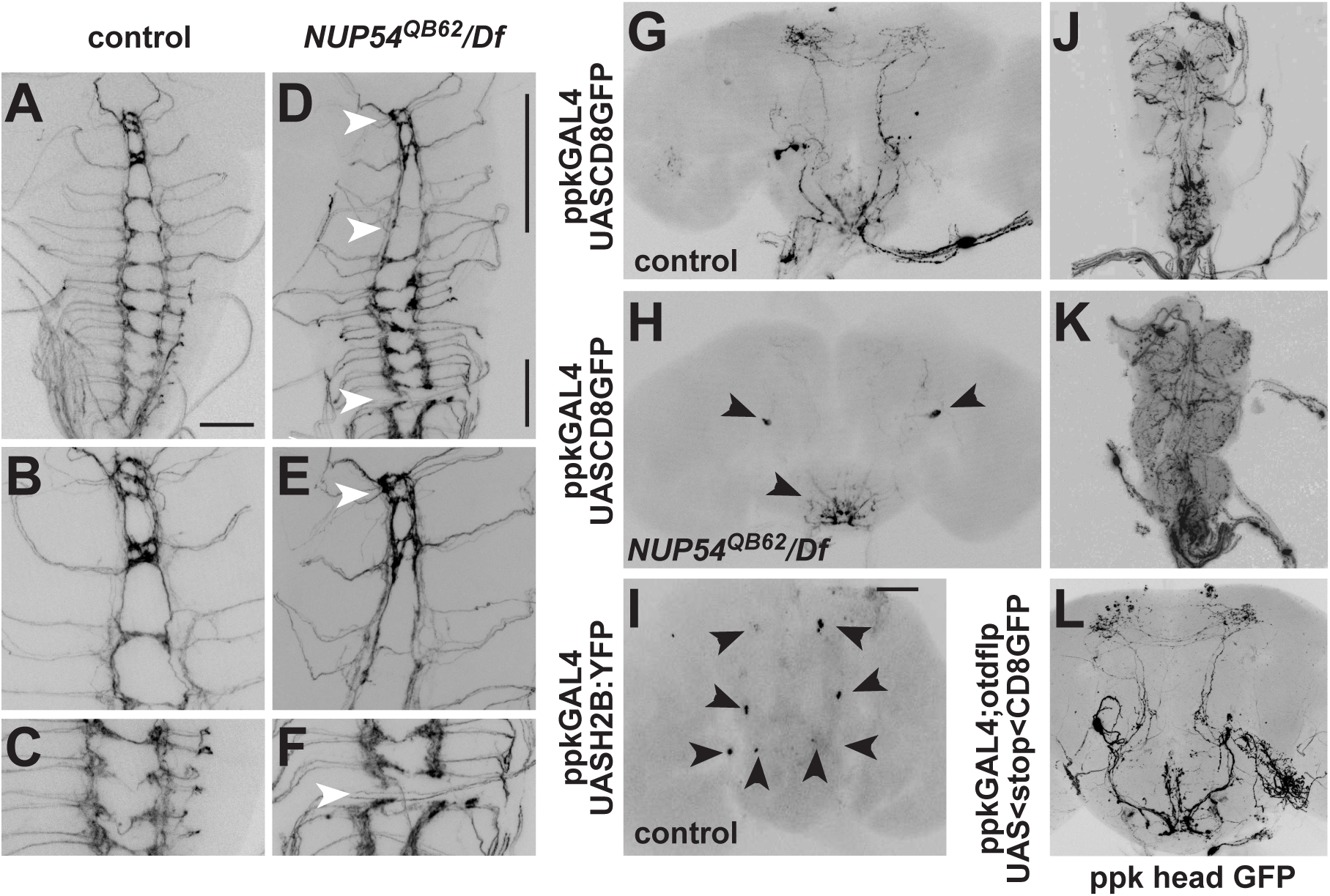
*Nup54* is required in *pickpocket* neurons to establish neuronal projections. **A-F)** Larval ventral nerve cord expressing CD8::GFP from *UAS* by *ppkGAL4* in wild type (A-C) and *Nup54*^*QB62*^*/Df(2R)9B4*. Magnified corresponding areas indicated in (D) are shown for wild type (B and C) and *Nup54*^*QB62*^*/Df(2R)9B4* (E and F). Arrowheads indicate missing commissures or connectives. **G, H)** Adult brain expressing CD8::GFP from *UAS* by *ppkGAL4* in wild type (G) and *Nup54*^*QB62*^*/Df(2R)9B4* (H). Arrowheads point towards cell bodies of *ppk* expressing neurons with absent neuronal arborizations. **I)** Adult brain expressing histon2B::YFP from *UAS* by *ppkGAL4* in wild type. Arrowheads point towards cell bodies of the four paired *ppk* expressing neurons in the adult brain. The scale bar in A is 50 µm and in H is 100 µm. **J, K)** Ventral nerve cord expressing CD8::GFP from *UAS* by *ppkGAL4* in wild type (J) and *Nup54*^*QB62*^*/Df(2R)9B4* (K). **L)** Adult brain expressing CD8::GFP only in head *ppk* neurons by intersection of *ppkGAL4 UAS FRTstopFRT CD8GFP* expression with *otdflp*, which expresses only in the brain.

So far it has been shown that *ppk* expresses in sensory neurons projecting to the brain [23-26]. A closer examination of *ppk* expression in the adult female brain, however, revealed four paired neurons consistently expressing a nuclear histon2B::YFP marker expressed from *UAS* with *ppkGAL4*. Strikingly, these neurons lacked projections in *Nup54*^*QB62*^*/Df(2R)9B4* females (Figure 2G-I). Since ascending neurons from the genital tract and the ventral nerve cord project to the brain, we used *flipase (flp)* expression under an *orthodenticle* (*otd*) promoter [39], which is restricted to the brain to determine whether the arborizations in the brain are from these four *ppk* neuron-pairs in the brain. No difference in the projections in the central brain was observed when *ppk* expression was restricted to the brain (Figure 2L) with an intersectional approach (Figure 3A) and accordingly, expression genital tract ppk neurons was absent (Figure 3 B, C).

**Figure 3:**
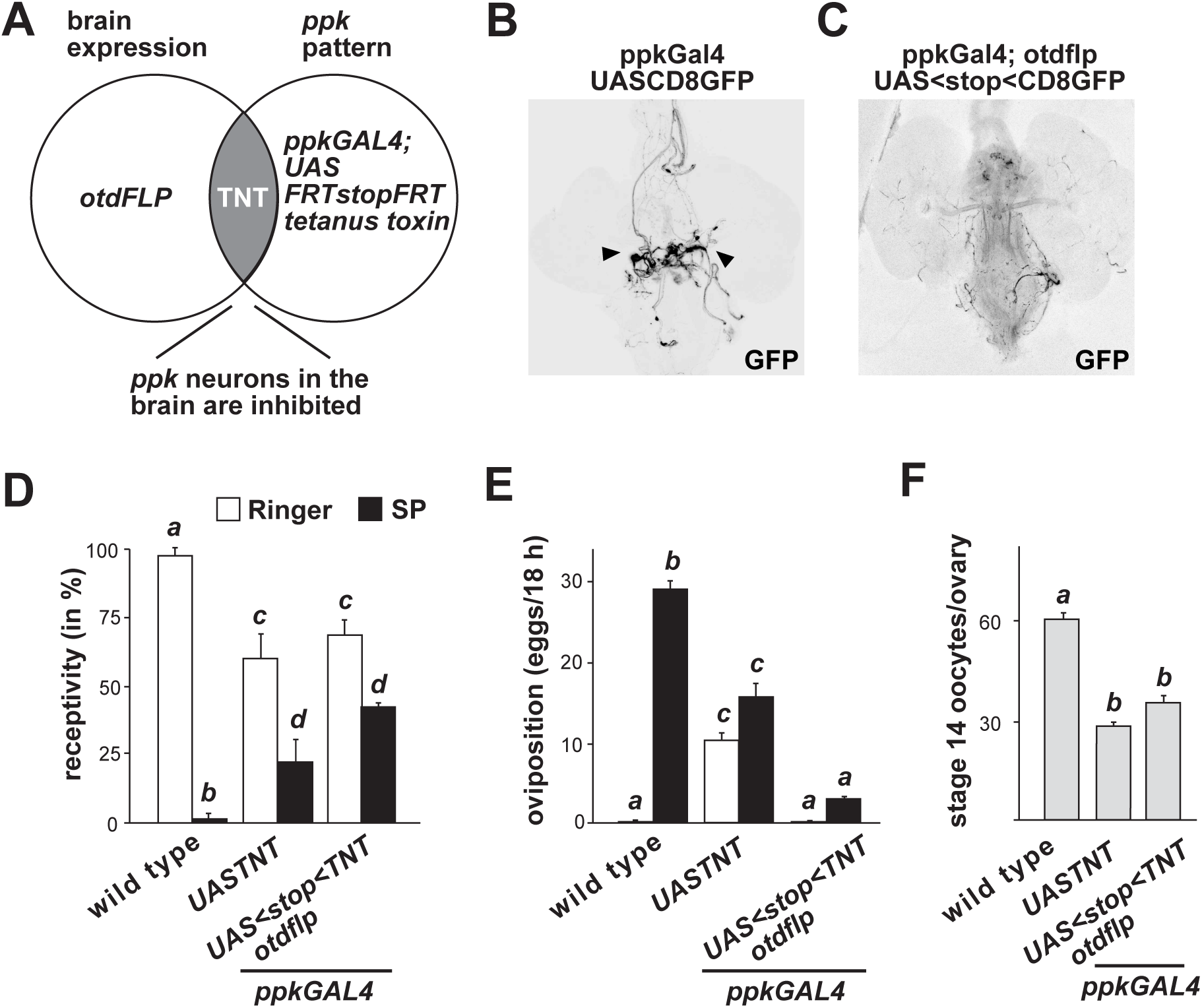
Brain *pickpocket* neurons are part of the circuit directing egg laying. **A)** Schematic of the intersectional gene expression approach to direct expression of mSP to brain *ppk* expressing neurons using brain expressed *otdflp* and *UAS FRTGFPstopFRTTNT*. **B, C)** Intersectional directed expression of *ppk* to the head in *ppkGAL4/UAS FRTGFPstopFRTCD8GFP; otdflp* removes expression in genital tract neurons. **D)** Receptivity of control and females expressing TNT in all (*ppkGAL4*) or only brain *ppk* neurons (*ppkGAL4 UAS stop TNT otdflp*) after sex-peptide (SP, black) or Ringer’s (R, white) injection measured by counting mating females in a 1 h time period 3 h after SP or R injection, respectively. Means with the standard error for three experiments with 10-24 females each are shown, and statistically significant differences are indicated by different letters (p≤0.05). **E)** Oviposition of control and females expressing TNT in all (*ppkGAL4*) or only brain *ppk* neurons (*ppkGAL4 UAS stop TNT otdflp*) after sex-peptide (SP, black) or Ringer’s (R, white) injection shown as means of eggs laid in 18 h with the standard error for 12-30 females each, respectively. Statistically significant differences are indicated by different letters (p≤0.001). **F)** Storage of stage 14 oocytes is not affected in virgin females expressing TNT in all (*ppkGAL4*) or only brain *ppk* neurons (*ppkGAL4 UAS stop TNT otdflp*). Means with the standard error from six ovaries each are shown, and statistically significant differences are indicated by different letters.

### *ppk* neurons in the brain are part of the circuit required for egg laying

To test whether *ppk* expressing neurons in the brain are involved in the post-mating response we used an intersectional gene expression approach based on brain-specific expression of *otd* to specifically inhibit neuronal transmission in brain *ppk* neurons by expression of tetanus-toxin (TNT) [39]. To achieve restricted expression of TNT in *ppk* brain neurons, we crossed *otdflp* flies with *ppkGAL4, UAS FRTGFPstopFRT TNT* flies and analysed the female progeny for PMRs (Figure 3A).

These experiments revealed that females with inhibited *ppk* neurons in the brain displayed a compromised response to SP as they did not fully reduce receptivity and did not lay the eggs stored in the ovaries after injection of SP (Figure 3B-D). In contrast, inhibition of all *ppk* neurons partially reduced receptivity in SP injected females. In addition, these females display increased egg laying as virgins, but SP does not further enhance egg laying (Figure 3B-D). Control females of *UAS TNT* without a *GAL4* driver showed a normal response to SP (data not shown).

Taken together, these results demonstrate an essential role for *ppk* neurons in the brain in inducing egg laying, and a partial requirement for reducing receptivity in response to SP.

### Channel Nups have a role in sex determination and dosage compensation

To further test the role of *Nup54* in the post-mating response we used *UASRNAi* knock-down of *Nup54* in either all neurons using *elavGAL4*^*C155*,^ or in subsets of neurons using *ppkGAL4, fruGAL4* and *dsxGAL4*. RNAi knock-down of *Nup54* with *elavGAL4* ^*C155*^, *ppkGAL4* and *fruGAL4* did not affect receptivity, or egg laying after SP injection further demonstrating that *Nup54* is required early in neuronal development (Figure S3A and B). In contrast, RNAi knock-down of *Nup54* with *dsxGAL4*, oviposition was not induced upon SP injection while receptivity was reduced normally (Figure S3A and B).

When knocking down *Nup54* with RNAi using *dsxGAL4*, we noticed that females laid eggs that did not develop. In addition, male survival was low (20%, n=107, *P{TRIP*.*HMC04733}*) or males were completely absent (n=107, *P{GD14041}v42153; P{GD14041}v42154*), indicating that *Nup54* has additional roles in dosage compensation as all mutants causing male lethality identified in excessive screens have been mapped to genes regulating this process [40, 41]. Therefore, we hypothesized that the other two channel Nups, Nup58 and Nup62, that form a complex with Nup54, also have roles in specifying the post-mating response and/or in sexual differentiation and dosage compensation. RNAi knockdown of *Nup58* (*P{TRIP*.*HMC05104}*) with *dsxGAL4* resulted in lethality with only few very weak female escapers. These females, however, were sterile due to the lack of ovaries and deformed genitals. Similar results were obtained by RNAi knock-down of *Nup62* with *dsxGAL4* resulting in male lethality (9%, n=81 and 2%, n=45 escapers with *P{TRIP*.*GLV21060}* and *P{TRIP*.*HMC03668}Since*) and females without ovaries (100%, n=74 and 43%, n=35 with *P{TRIP*.*GLV21060}* and *P{TRIP*.*HMC03668}*). In addition, these females also had deformed genitals and males displayed underdeveloped sex combs indicating a role for channel Nups in sexual differentiation (Figure S3C-F). Consistent with a regulatory role in development, the channel Nups display dynamic expression during development and in different tissues (Figure S4A-F).

### The role of insertion/deletion (indels) during the evolution of Nup54 function

Adaptation to new situations drives the evolution of hybrid incompatibility genes *Nup96* and *Nup160* [31-34]. Here, we used the combined analysis of polymorphism within *D. melanogaster* and divergence between *D. melanogaster* and *D. simulans* within the coding region of *Nup54* to test for deviations from expectations under the neutral model of molecular evolution. We found no deviation from the null neutral model hypothesis, as the ratio of nonsynonymous to synonymous polymorphisms did not significantly differed from the ratio of nonsynonymous to synonymous fixed differences between species (MK test: χ^2^= 0.09; P= 0.923). The test was also non-significant when we partition the analysis by exons. Also, the upstream noncoding region did not significantly depart from neutrality (Fisher’s exact test P= 0.135), but there was a trend towards an excess of upstream noncoding relative to silent (synonymous and intron) substitutions between species (dNC/dS=3.1) relative to polymorphisms within *D. melanogaster* (PNC/PS=2.0) (neutrality index: NI = 0.65).

Indels are a common form of genetic variation, being able to affect gene function and their pattern of evolution being subjected to selective forces [42-44]. We noticed that a stretch of amino acids of increasing length in more distantly related species is inserted in the FG repeat region (Figure S5A). We therefore hypothesized that this indel could alter the function of Nup54 in specifying the post-mating response. To test this hypothesis, we replaced the FG-repeat region from *D. melanogaster* with the region from *D. elegans* in the genomic rescue construct *gNup54ele* and tested its capacity to rescue viability of *Df(2R)9B4* and the post-mating response defects. Both the *D. melanogaster gNup54* and the chimeric *D. elegans* construct *gNup54ele* rescued viability of *Df(2R)9B4* to 91% (n=242) and 95% (n=381) without sex-bias, respectively. After injection of SP, females rescued with the *gNup54* or the *gNup54ele* construct showed normal post-mating responses in receptivity and oviposition arguing that this indel is not the source for an altered post-mating response (Figure S5B and C).

In the *Nup54* promoter region rapid evolution is identifiable by the presence of four hotspots for substitutions between species, all within 1 kb upstream of the *Nup54* transcription start site (Figure 4A and B). Interestingly, one substitutional hotspot (−359 to −295) includes an indel that overlaps with the location of the characterized *Nup54*^*QB62*^ regulatory allele (Figure 4B). The deletion in the *Nup54*^*QB62*^ allele in fact is to a large extent a wild type condition in sister species (Figure 4A). To test whether the alteration in the promoter of sister species impact on Nup54 expression, we replaced the *D. melanogaster* promoter in *gNUP54* construct with the promoter of *D. simulans* to generate *gNUP54*^*Dsim*^. Indeed, expression in salivary glands was reduced and we noted changes in the brain (Figure S1 A, B compared to S1 E, F). While it is unclear whether these changes in the promoter sequences were been driven by selection, it has clearly evolved in the context of the post-mating response to SP.

**Figure 4:**
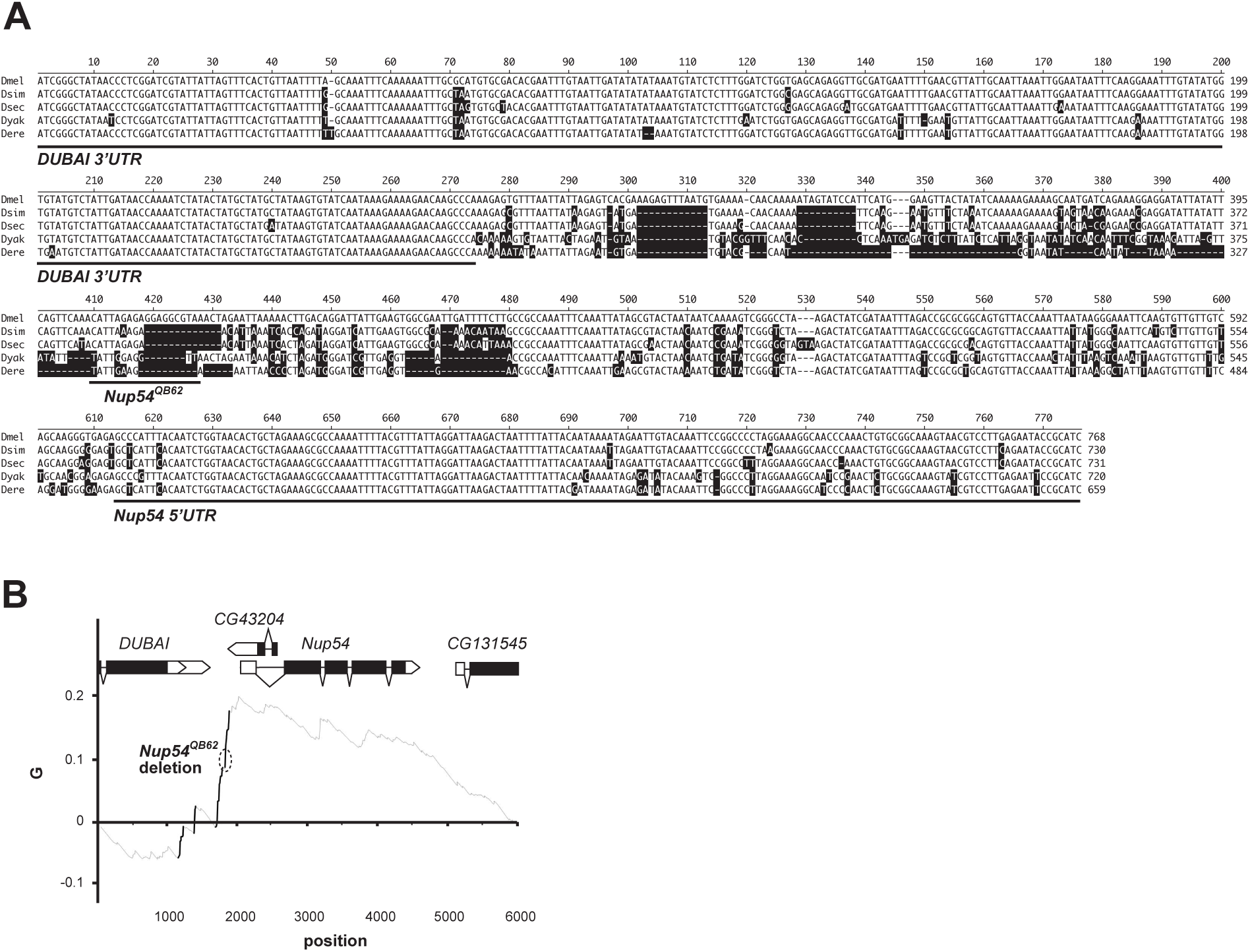
Nup54 promoter region is diverged in closely related species. A) Sequence alignment of the *Nup54* promoter region from closely related species. Nucleic acids deviating from *D. melanogaster* are indicated in black. Transcribed parts of the *DUBAI* 3’UTR and the *Nup54* 5’UTR, as well as the *Nup54*^*QB62*^ allele are underlined. B) Plot of cumulative differences along the sequence (G) between the relative occurrences of nucleotide changes and their position from the alignment of the gene region around *Nup54* between *D. melanogaster* and *D. simulans*. Positions in the alignment with significant stretches of substitutions (hotspots) are identified by solid lines. The deletion in the *Nup54*^*QB62*^ allele indicated by a circle with a dashed line.

## Discussion

Here we show that Nup54 which is part of the central transport channel of the nuclear pore exerts a role in specifying the neuronal circuits involved in mediating the SP induced post-mating switch. Viable alleles of these genes display an egg retainer phenotype and are insensitive to SP with regard to reducing receptivity. Nup54 is required for wiring of *ppk* expressing neurons in the brain that constitute part of the circuitry required for egg laying. The role of Nup54 and the other two channel Nups, Nup58 and Nup62, in sexual differentiation, however, seems to be more fundamental as reducing their expression levels impacts on general sexual differentiation of external and internal sexual features, such as female genitals and ovaries, but also on dosage compensation in males.

## Roles for Nups in development

A key role for Nups in sexual differentiation seems rather unexpected as the function of channel Nups in directing constitutive export of mRNAs and proteins from the nucleus to the cytoplasm has been viewed as static for a long time. However, a number of recent studies have brought about developmental and neuronal roles for the nuclear pore and nucleocytoplasmic transport reflected in a number of human diseases including neurodegeneration associated with specific Nups [45-47]. A central aspect of these more profound roles for Nups lay in their varying expression levels between different cell types, tissues and in development. Nup210’s key role in muscle and neuronal differentiation are associated with tissue-specific expression [48]. Likewise, high expression levels of Nup153 are critical for maintenance of ESC pluripotency as reducing levels results in neuronal differentiation [49]. Also, channel Nup62 shows increased expression in various epithelia and is required for proliferation [50]. In *Drosophila*, channel Nups 54, 58 and 62 show increased expression in the larval brain and adult gonads [51], but a more specific role is indicated in sexual development in *Drosophila* from broad RNAi knock-down with *dsxGAL*, recapitulating *dsx* expression cell-autonomously directing sex determination [40, 41]. Our findings that channel Nups have a role in sexual differentiation add to this view that the nuclear pore has key roles in differentiation and likely also neuronal function.

### Specification of neuronal circuits for receptivity and egg laying are separable

It has previously been argued that the neurons co-expressing *fru, dsx* and *ppk* in the genital tract sense SP and direct the SP induced post-mating [24-26]. However, our data from expressing a *Nup54* rescue construct in various cell types suggests a more complex picture including separation of the paths leading to reduced receptivity and increased egg laying. Here, expression of Nup54 in *fru* and *dsx* neurons, but not *ppk* neurons rescues receptivity while egg laying is not rescued in all three expression patterns. Although this effect could be explained by different expression levels of *fru, dsx* and *ppk* GAL4 lines, this is not observed and they all express strongly. Consistent with a requirement for Nup54 early in development, *dsx* is expressed earlier than *fru* and *ppk*. Differential effects on receptivity, but not egg laying by *dsx* driven rescue by Nup54, however, more likely indicates that receptivity and egg laying are governed by different neuronal circuits. These results are supported by previous observations, that G(o) is not required in *fru*, but needed in *dsx* and *ppk* neurons to reduce receptivity [9]. Likewise, membrane-tethered SP can only induce oviposition in the absence of SPR in *dsx*, but not *fru* and *ppk* neurons [9].

### Roles of Nups in speciation

New species can arise through selection of new features enhancing display of sexual attributes and altered courtship behavior. A driving force to speciation can be sexual conflict imposed by male-directed post-mating responses by females [28-30]. In *Drosophila melanogaster*, females respond to male-derived SP by reducing receptivity and increasing egg laying, but need to adapt their physiological status to environmental conditions. Thus, the possibility to limit the male influence imposed by SP is in the female interest of optimizing reproductive success when resources to produce eggs are scarce. In this context, Nup54 regulation could impact on egg laying in dedicated circuits such as brain *ppk* neurons. Consistent with this interpretation, the Nup54 promoter region has undergone a dramatic change compared to closely related species, while no adaptive changes were detected in the protein coding part. Likewise, exchanging the FG repeat of Nup54 mediating nucleo-cytoplasmic transport, which shows most variability between *D. melanogaster, D. elegans* and other closely related species had little impact on the post-mating response. Since Nup54 is required in all cells, varying its concentration could have profound effects on general expression of genes and select for a specific compensatory genetic element from the mating partner to prevent deleterious effects on the fitness of the progeny[31-34]. Such interpretation is well in line with roles of essential Nup96 and Nup160 in speciation as they have been identified in causing hybrid sterility or lethality. Intriguingly however, Nup96 and Nup160 are part of the larger NUP107 subcomplex and four out of eight proteins from this complex are under adaptive selection in *D. simulans* [32]. The driving forces behind the role of Nups in speciation and the molecular mechanism leading to speciation, however, need to be further explored. Intriguingly, a role for *fru* in female receptivity has been mapped in interspecies hybrids to the sex-specific transcript [52]. Since the female isoform seems only to be translated before the adult stage, expression of the female isoform might be linked to its long 5’UTR that could impact on nuclear-cytoplasmic export.

Here we have identified a few *ppk* expressing neurons in the brain as a cellular focus preventing female escape from male manipulation. Changes in neuronal wiring likely directed in response to sexual conflict arising from male-derived SP to direct the female post-mating response marks an early event in the splitting of species and links differentiation of key neurons involved in female control of reproduction to fitness as a result of sexual conflict. Our results indicate a central role for the nuclear pore in implementing alterations of gene expression impacting on neuronal wiring at the onset of speciation processes.

## Materials and Methods

Flies were kept on standard cornmeal-agar food (1%industrial-grade agar, 2.1% dried yeast, 8.6% dextrose, 9.7% cornmeal and 0.25% Nipagin, all in (w/v)) in a 12 h light: 12 h dark cycle. Sexually mature 3–5 day-old virgin females were injected and examined for their post-mating behaviors as described previously [12, 22]. For injections, virgin female flies were cooled to 4° C and 3 pmol SP in 50 nl Ringer’s solution was injected. Ovaries were analyzed as previously described [12]. ANOVA followed by planned pairwise comparisons with Fisher’s protected least significant difference was done for statistical analysis using STATVIEW or Graphpad prism.

The *UAS* construct with a N-terminally HA-tagged Nup54 was generated in a three way ligation with fragments of *Nup54* amplified from cDNA by RT-PCR that were cloned into a modified *pUC, pUC 3GLA UAS HA*, with NheI and Acc65I with a blunt site in between. The 5’ part was amplified using primers 8831F1 (CATCGCTAGCGCCTGCAGGATCGTTCTTCGGATCCAACACGTCGCTGG) and 8831R1 (CTGAGAGTTCTCTGCAGAAGTTAAGAGCCAC), and the 3’ part with primers 8831F2 (CTGTCAAGCCACACCAGCAACAAGTGATTC) and 8831R2 (GACAGGTACCTATCACGATTGTCGCAGCTCGGGCAGTC) by PCR with Pwo (Roche) and sequenced. The genomic rescue construct was generated in a three way ligation by cloning the promoter fragment amplified from genomic DNA with primers 8831F1g (GTGGAATTCCGGAGGCCACTAGAACATATACTTGTC) and 8831R1g (GGCGTGCTTGTTGCTCCCAGCGACGTGTTG) and the cDNA part amplified from the *UAS* construct with primers 8831F3 (GGCCAAAACAACCGGTGGCCTCTTCGGATC) and 8831R3 (GGCGTGCTTGTTGCTCCCAGCGACGTGTTGTAGCTCGAGGATTGTCGCAGCTCGGGCAGTC) using EcoRI and XhoI sites into a modified *pUC, pUC 3GLA* that adds a C-terminal HA tag and sequencend [53]. Swapping the *NUP54* wild type promoter was done by digesting the parent plasmid with BspEI and NgoMIV to insert the *Nup54*^*QB62*^ promoter or the promoter from *Drosophila simulans* (S-23, Ethiopie 225, Welo Ataye River, F. Leumeunier, gift from S. Collier Cambridge) from a PCR amplified with primers 8831F3BspE (GCTTAGGATCCGATCGCGTGGAATTCCGGAGGCCACTAGAACATATACTTGTC) and 8831promR (CGAAGAGGCCACCGGTTGTTTTGGCCGGCGTGCTTGTTGCTCCCAGCGACGGTTG). The resulting *pUC 3GLA gNUP54*^*QB62*^ and *pUC 3GLA gNUP54*^*Dsim*^ constructs were validated by sequencing with primer 8831promR2seq (GGCTGGTTGCAGCTGTGCCTCCAAAC). The *D. elegans* rescue construct was generated in a three way ligation by replacing the part between NgoMIV and MfeI with the corresponding part amplified from *D. elegans* cDNA by RT-PCR with primers 8831Fele (CCAAAACAACcGGaGGCCTCTTCGGAAC) and 8831R1ele (CGTGGATCCGAAGGCTCCGCCCCCAAAGCCAGTG), and a BamHI/MfeI fragment obtained from the UAS construct. Transgenic flies were generated by phiC31-mediated transformation using landing site 76A (*PBac{y+ -attP-3B}VK00002*), where the GFP marker had been removed by Cre/Lox mediated recombination. *Df(2R)9B4* was generated by FLP/FRT-mediated recombination between two transposon insertion lines, *P{XP}CG8525*^*d06853*^ and *PBac{RB}DUBAI*^*e00699*^ as described [54, 55]. The *tubGal4* line was a third chromosomal insert (Bloomington #5138), *UAS FRTGFPstopFRT TNT* and the other lines have been previously described [9, 56, 57]. Antibody stainings were done as previously described [58, 59].

Genomic DNA was extracted form *Nup54*^*QB62*^ as described [60]. The promoter fragment was amplified with primers 8831pF1 (GGATCTGGTGAGCAGAGGTTGCGATG) and 8831pR1 (GCCGCACAGTTTGGGTTGCCTTTC) and sequenced. Reverse transcription quantitative polymerase chain reaction (RT-qPCR) was done by extracting total RNA using Tri-reagent (SIGMA) and RT was done with Superscript II (Invitrogen) according to the manufacturers instructions using an oligo dT primer. For qPCR 1.5 µl cDNA with the SensiFAST SYBR No-Rox kit (Bioline) was used with primers Nup54qF1 (CTGCCACAGCGAAGATACT) and Nup54qR1 (CAGCATGTTCTGTAGCTTGGTGC), and ewg4F1 and ewg5R1 [61]. Amplification was done in a Applied Biosystems ABI Prism 7000 with 3 min initial denaturation at 95° C followed by 40 cycles with 15 sec denaturation at 95°C and 60 sec extension at 60° C. Quantification was done according to the ΔCT method [62].

Structural analysis was done by PyMol. The *Nup54* ORF and extended gene sequences were retrieved from FlyBase (flybase.org) and aligned using muscle within MEGA (reference). *D. melanogaster* polymorphism data was retrieved from the Drosophila Genetic Reference Panel (http://dgrp2.gnets.ncsu.edu/) and used along with interspecies divergence data (*D. melanogaster – D. simulans*) to conduct McDonald Kreitman’s tests of selection for the ORF and extended gene region. Tests were conducted by comparing nonsynonymous to synonymous substitutions and polymorphisms (ORF) as well as noncoding upstream to silent (within gene synonymous and intron) substitutions and polymorphism [63-65].

We also tested for evidence of nonrandom accumulation of substitutions along the *Nup54* extended gene region. The method tests for significant deviations from a uniform distribution of substitutions using an empirical cumulative distribution function. The function (G) detects monotonic increases in substitutions (n) measured as the difference between the relative occurrence of a nucleotide change and its relative position in the alignment [66, 67]. Whether differences between the values of the G function (ΔG) between substitutional events deviates from a random accumulation of changes are tested using Monte Carlo simulations to produce 100,000 samples of n events by sampling sites without replacement along the alignment [66, 67].

## Acknowledgments

We thank T. Schüpbach, B. Dickson, S. Goodwin, C. Rezaval, D. Anderson, S. Collier, the Bloomington stock center, the Vienna Drosophila RNAi Center and the Arizona species stock center for fly lines and the University of Cambridge Department of Genetics Fly Facility for injections, N. Arora for help with behavioral assays, D. Scocchia for help with graphics and qPCR, and T. Dix for help with PyMol, C. Rezaval and J.C. Billeter for comments on the manuscript. We are indebted to Eric Kubli for his support when this study was initiated. This work was supported by the Biotechnology and Biological Science Research Council to MS and the Natural Sciences and Engineering Research Council of Canada to AC.

## Authors’ contribution

MS conceived and directed the project. IUH and MS performed molecular biology experiments, IUH, MN and MS performed genetic experiments, and AC performed sequence analysis. All authors analyzed data. MS wrote the manuscript with support from IUH and AC. All authors read and approved the final manuscript.

