## Supplemental Information for "Channel nuclear pore protein 54 directs sexual differentiation and neuronal wiring required for female reproductive behaviors in *Drosophila*"

#### Supplementary Figures

**Supplementary Figure S1: Nup54 expression is altered by the *QB62* deletion or the *Drosophila simulans* promoter without gross disorganization of the brain.**

**A-F)** Expression of Nup54 from a C-terminally HA tagged genomic rescue construct with the wild type promoter (*gNup54*), the promoter from the NupQB62 allele (*gNup54*$^{QB62}$) or *Drosophila simulans* (*gNup54*$^{Dsim}$) in salivary glands (A, C and E, top) and the adult brain (B, D, and F, top) above nuclear DAPI staining (middle) and the merged picture (bottom) with scale bars indicating 20 µm and 100 µm for salivary glands and brains, respectively.

**G-J)** Projections of *dsx* neurons visualized by expression of membrane-bound CD8GFP from *UAS* by *dsxGAL4* show no gross alterations in *Df(2R)9B4 gNup54*$^{QB62}$ compared to wild type. Changes in expression are indicated by arrowheads.

**Supplementary Figure S2: *Nup54* is required before neuronal maturation for establishing the post-mating response**

**A)** Receptivity of control (blue) and transheterozygous *Nup54*$^{MB03363}$*/Df(2R)9B4* expressing *UASNup54::HA* with *tubGAL4* (purple) ubiquitously, in neurons with *elavGAL4*$^{C155}$ (yellow) and in *ppkGAL4* (orange), *fruGAL4* (green) and *dsxGAL4* (red) patterns after sex-peptide (SP, dark color) or Ringer's (R, light color) injection measured by counting mating females in a 1 h time period 3 h after SP or R injection, respectively. Means with the standard error for three experiments with 18-21 females each are shown, and statistically significant differences are indicated by different letters (p≤0.001).

**B)** Oviposition of control (blue) and transheterozygous $Nup54^{MB03363}/Df(2R)9B4$ expressing *UASNup54::HA* with *tubGAL4* (purple) ubiquitously, in neurons with $elavGAL4^{C155}$ (yellow) and in *ppkGAL4* (orange), *fruGAL4* (green) and *dsxGAL4* (red) patterns after sex-peptide (SP, dark color) or Ringer's (R, light color) injection shown as means of eggs laid in 18 h with the standard error for 10-16 females each, respectively, and statistically significant differences are indicated by different letters (p≤0.001).

**Supplementary Figure S3: *Nup54 RNAi* in *doublesex* expressing neurons reveals a separable sex-peptide response in receptivity and oviposition and a role in sexual differentiation.**

**A)** Receptivity after *Nup54 RNAi* knock-down from *UAS P{GD14041}v42153; P{GD14041}v42154* inserts in neurons with $elavGAL4^{C155}$ (yellow) and in *ppkGAL4* (orange), *fruGAL4* (green) and *dsxGAL4* (red) patterns after sex-peptide (SP, dark color) or Ringer's (R, light color) injection measured by counting mating females in a 1 h time period 3 h after SP or R injection, respectively. Means with the standard error for three experiments with 16 females each are shown, and statistically significant differences are indicated by different letters (p≤0.001).

**B)** Oviposition after *Nup54 RNAi* knock-down from *UAS P{GD14041}v42153; P{GD14041}v42154* inserts in neurons with $elavGAL4^{C155}$ (yellow) and in *ppkGAL4* (orange), *fruGAL4* (green) and *dsxGAL4* (red) patterns after sex-peptide (SP, dark color) or Ringer's (R, light color) injection shown as means of eggs laid in 18 h with the standard error for 8 females each, respectively, and statistically significant differences are indicated by different letters ($p \leq 0.001$).

**C, D)** Genitals of control females (C) and females expressing *Nup62 RNAi* from *UAS* with *dsxGAL4* (D). The scale bar in B is 20 µm.

**E, F)** Front legs of control males (E) and males expressing *Nup62 RNAi* from *UAS* with *dsxGAL4* (F). Arrowheads indicate the position sex combs. The scale bar in B is 100 µm.

**Supplementary Figure S4: Expression of *Nup54*, *Nup58* and *Nup62*.**

**A-C)** Profile of *Nup54*, *Nup58* and *Nup62* expression during development from RNA seq.

**D-F)** Profile of *Nup54*, *Nup58* and *Nup62* expression in various tissues from microarrays.

**Supplementary Figure S5: Nup54 is highly conserved, but variation in the FG repeat region is not the cause of an altered post-mating response.**

**A)** Sequence alignment of Nup54 from closely related species. Amino acids deviating from *D. melanogaster* are indicated in black. Intron positions are indicated by black arrowheads and the stop codon of the *Nup54$^{MB03363}$* allele is indicated by a red arrow head. Green and white filled arrowheads indicate maintained (>1%) and rare (<1%) polymorphisms with amino acid changes indicated on top. The line below the sequence indicates the NgoMIV-BamHI fragment that was replaced in the *gNup54elegans* construct. Nucleotides 1-115 according to human Nup54 impasse the FG region, nucleotides 116-346 the α/β region and nucleotides 346-494 the α-helical region. The amino acids in exon 3 have been shown to bind to Nup62 and the amino acids in exon 5 bind to Nup58.

**B)** Receptivity of wild type, *gNup54* and *gNup54elegans* females homozygous for *Df(2R)9B4* after sex-peptide (SP) or Ringer's (R) injection measured by counting mating females in a 1 hr time period 3 hr after SP or R injection, respectively. Means with the standard error for three experiments with 8-15 females each are shown, and statistically significant differences are indicated by different letters.

**C)** Oviposition of *gNup54,* and *gNup54elegans* females homozygous for *Df(2R)9B4* after sex-peptide (SP) or Ringer's (R) injection shown as means of eggs laid in 18 h with the standard error for 10 females each, respectively. Statistically significant differences are indicated by different letters.

### Figure S1
### Nallasivan et al

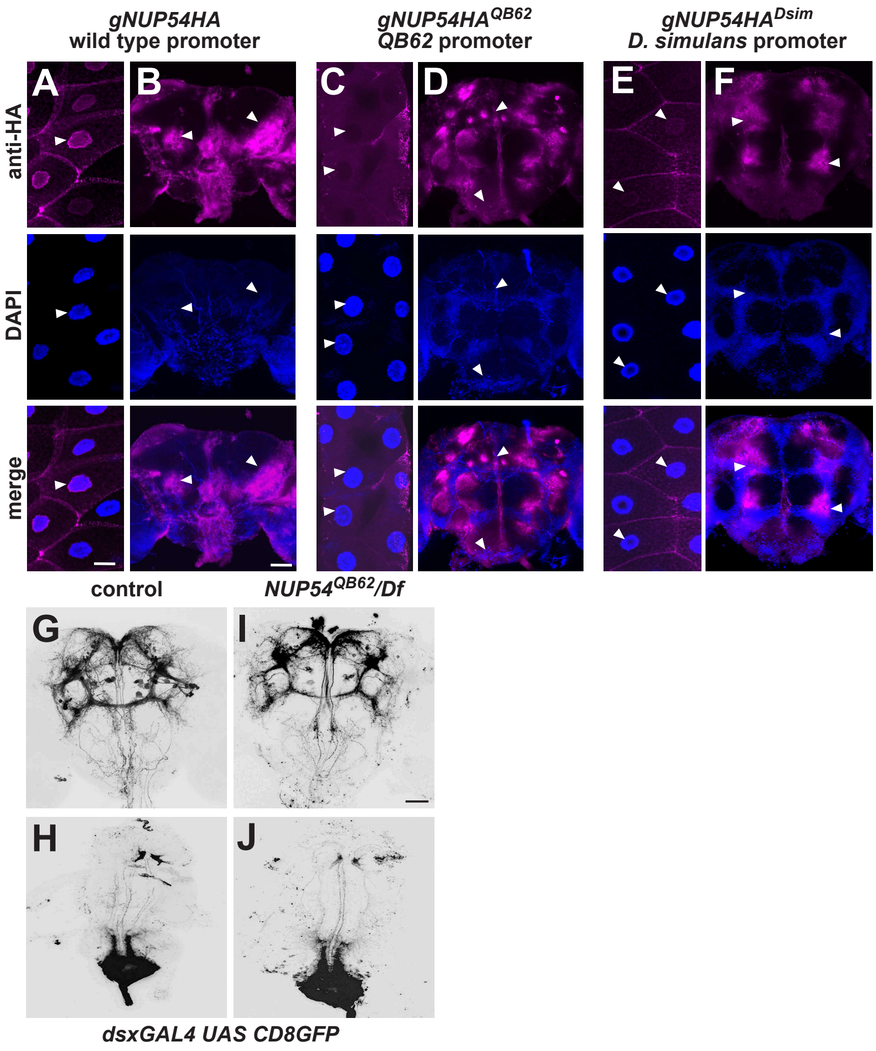

### Figure S2
### Nallasivan et al

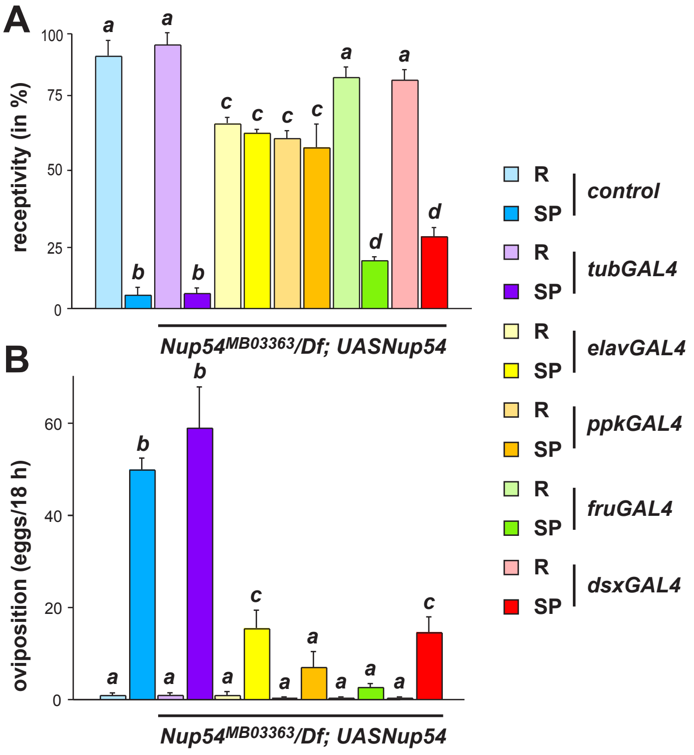

### Figure S3
### Nallasivan et al

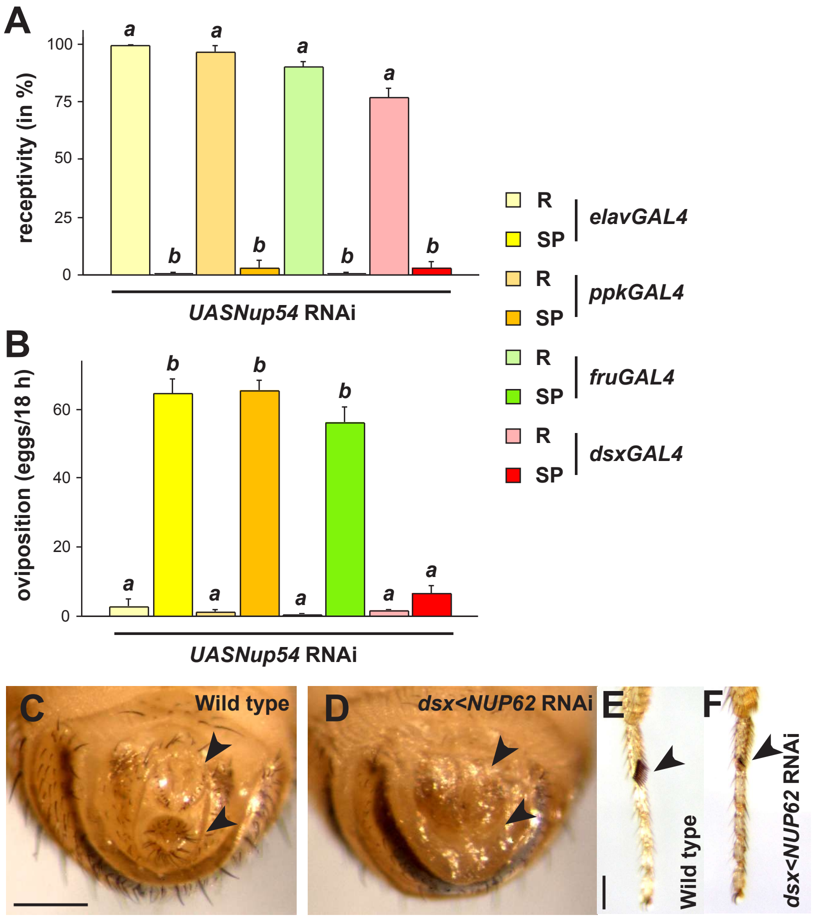

### Figure S4
### Nallasivan et al

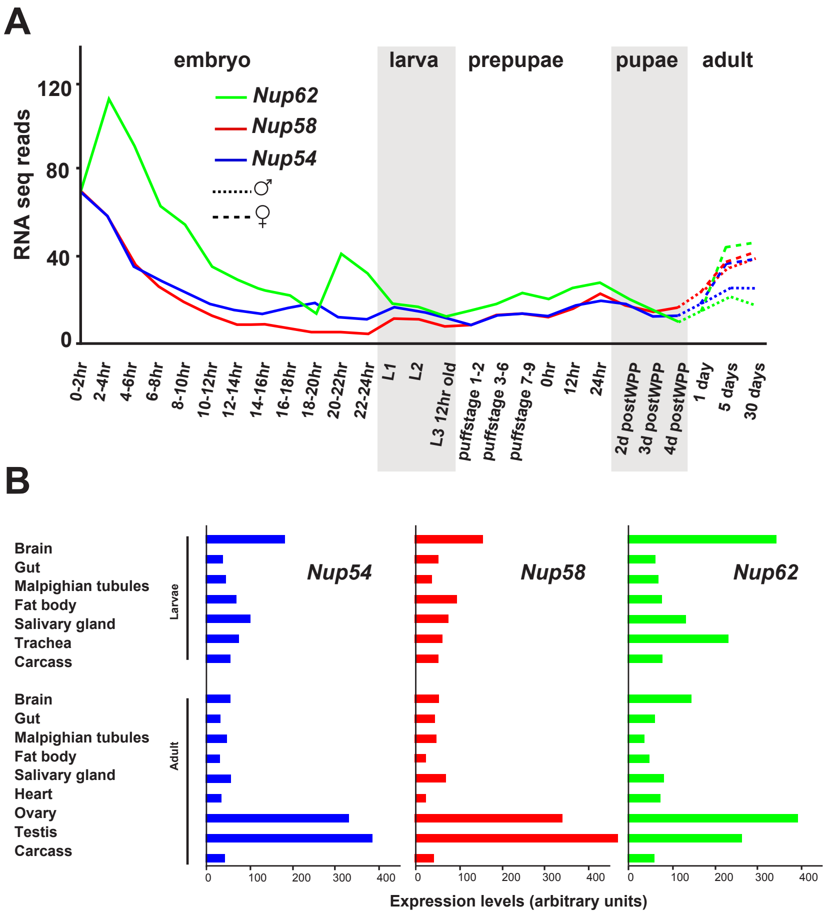

### Figure S5
### Nallasivan et al

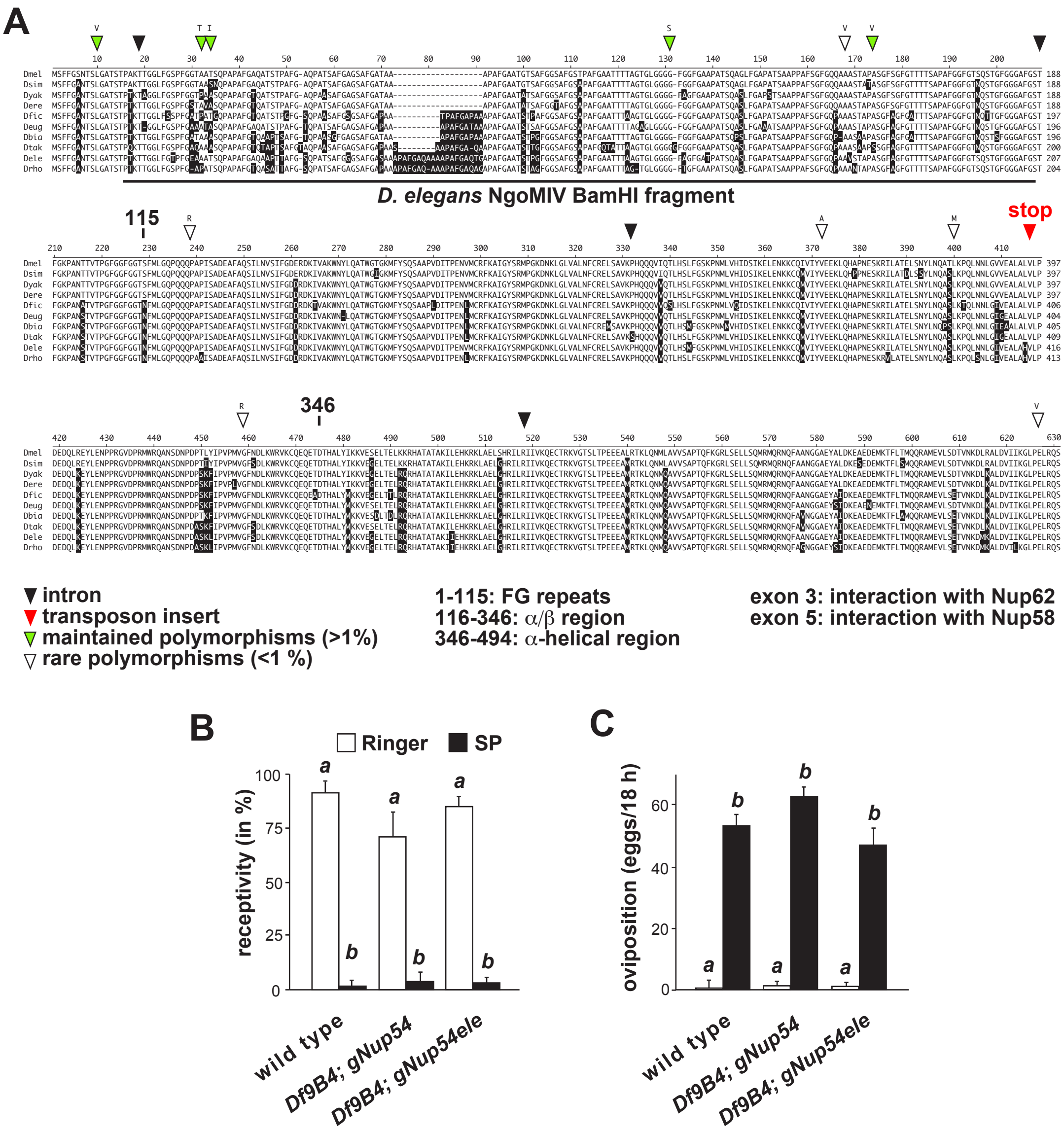
